# Transcriptional reprogramming and constitutive PD-L1 expression in melanoma are associated with dedifferentiation and activation of interferon and tumor necrosis factor signaling pathways

**DOI:** 10.1101/2021.06.15.448594

**Authors:** Antonio Ahn, Euan J. Rodger, Gregory Gimenez, Peter A. Stockwell, Matthew Parry, Peter Hersey, Aniruddha Chatterjee, Michael R. Eccles

## Abstract

Melanoma is the most aggressive type of skin cancer, with increasing incidence worldwide. Advances in targeted therapy and immunotherapy have improved the survival of melanoma patients experiencing recurrent disease, but unfortunately treatment resistance frequently reduces patient survival. Resistance to targeted therapy is associated with transcriptomic changes, and has also been shown to be accompanied by increased expression of programmed death ligand 1 (PD-L1), a potent inhibitor of immune response. Intrinsic upregulation of PD-L1 is associated with genome-wide DNA hypomethylation and widespread alterations in gene expression in melanoma cell lines. However, an in-depth analysis of the transcriptomic landscape of melanoma cells with intrinsically upregulated PD-L1 expression is lacking. To determine the transcriptomic landscape of intrinsically upregulated PD-L1 expression in melanoma, we investigated transcriptomes in melanomas with constitutive versus inducible PD-L1 expression (referred to as PD-L1_CON_ and PD-L1_IND_). RNA-Seq analysis was performed on seven PD-L1_CON_ melanoma cell lines and ten melanoma cell lines with low inducible PD-L1_IND_ expression. We observed that PD-L1_CON_ melanoma cells had a reprogrammed transcriptome with a characteristic pattern of dedifferentiated gene expression, together with active interferon (IFN) and tumor necrosis factor (TNF) signalling pathways. Furthermore, we identified key transcription factors that were also differentially expressed in PD-L1_CON_ versus PD-L1_IND_ melanoma cell lines. Overall, our studies describe transcriptomic reprogramming of melanomas with PD-L1_CON_ expression.

**Simple Summary:** Melanoma, an aggressive form of skin cancer, is frequently associated with drug resistance in the advanced stages. For instance, frequently resistance is observed to sequential treatment of melanoma with targeted therapy and immunotherapy. In this research, the authors investigated whether potential transcriptional mechanisms and pathways associated with PD-L1 protein expression could underlie targeted therapy drug resistance in melanoma. The authors found a PD-L1 expression transcriptional pattern underlies resistance to targeted therapy in a subgroup of melanomas. These melanomas were markedly dedifferentiated, as compared to melanomas that were not drug resistant. Understanding changes in transcription and molecular pathways that lead to drug resistance could allow researchers to develop interventions to prevent drug resistance from occurring in melanoma, which could also be relevant to other cancer types.

## 1. Introduction

Melanoma is the most deadly form skin cancer, as it frequently presents with highly aggressive features, including high propensity to metastasise and innate drug resistance. Moreover, these features frequently occur at a relatively early stage in the growth of the tumor [1]. Treatment of metastatic melanoma has been revolutionized over the last decade, as greater understanding has emerged of two critical hallmarks of melanoma: Firstly, a large proportion of melanomas (50-65%) are addicted to MAPK signalling through *BRAF* or *NRAS* mutations. In keeping with this, inhibition of the oncogenic BRAF protein has resulted in significant response rates in *BRAF* mutant melanomas. Secondly, irrespective of mutation status, melanoma is frequently dependent on immune suppression through programmed death 1 (PD1) signalling, upon the binding of ligand, either PD-L1 or PD-L2 [2]. Anti-PD1 therapy, which inhibits binding of PD-L1 or PD-L2 to the PD1 receptor, reactivates immune responses and has greatly improved melanoma patient survival [3,4]. Unfortunately, for both of these therapies, resistance inevitably develops, and currently no robust biomarker has been identified that is able to predict patient response. Neither PD1 nor PD-L1/PD-L2 expression accurately predict response, and the basis for intrinsic resistance to anti-PD1 treatment of melanoma is incompletely understood. Nevertheless, PD-L1 is constitutively expressed in some melanomas despite the absence of immune cell infiltration in the tumor. Furthermore, PD-L1 has also been shown to be upregulated upon development of resistance to MAPK pathway inhibitors, and is accompanied by tumor cell-intrinsic transcriptomic reprogramming [5,6]. However, it is currently unclear how intrinsic transcriptomic reprogramming occurs.

As we have previously described [7,8] PD-L1 expression in melanoma can broadly be categorised into mechanisms that are mediated by the presence or absence of tumor infiltrating lymphocytes (TILs). In the presence of TILs, tumor-associated PD-L1 expression is predominantly induced via interferons (IFN) [9-11] and/or cytokines, such as tumor necrosis factor (TNF) [12] secreted from TILs. We refer to melanoma cells with inducible PD-L1 expression as PD-L1_IND_. In contrast, PD-L1 expression without TILs, is largely mediated cell-intrinsically (or constitutively) via genetic or epigenetic mechanisms, and we refer to these type of melanoma cells as PD-L1_CON_ [2,13].

In this study, we describe transcriptomic features of melanoma cell lines with constitutive high expression of PD-L1 (PD-L1_CON_) compared to melanoma cell lines with low inducible levels of PD-L1 expression (PD-L1_IND_). We found that PD-L1_CON_ expression was associated with a reprogrammed transcriptomic state, inclusive of dedifferentiation, and active innate inflammatory pathways associated with IFN and TNF signalling and reduced oxidative phosphorylation. Overall, changes in the expression of key transcription factors were observed that drive dedifferentiation (such as the loss of *MITF* and *SOX10*), as well as key transcription factors that enhance IFN and TNF signalling and PD-L1 expression (such as *IRF1*). We additionally found that the altered expression of transcription factors and transcriptional reprogramming correlated with the PD-L1_CON_ mRNA expression, and with factors associated with resistance to MAPK pathway inhibitors.

## 2. Materials and Methods

### Selection and culture of melanoma cell lines

We analysed transcriptome feature of seven PD-L1_CON_ lines, which includes four (CM143.pre, CM143.post, NZM9, NZM40) lines from our previous study [7] and three additional new PD-L1_CON_ cell lines (MM127, MM595 and COLO239F) that were included in this study. These three cell lines were kindly provided by professor Glen Boyle from the QIMR Berghofer Medical Research Institute. For PD-L1_IND_ group, we have analysed 10 cell lines, which includes six (CM138, CM150.post, CM145.pre, CM145.post, NZM22, NZM42) lines from our previous study [7] and four new cell lines (NZM12, NZM15, WM115 and WM2664). The CM138, CM145.pre, CM145.post, CM150.post, CM143.pre and CM143.post lines were cultured in DMEM medium (Invitrogen) supplemented with 10% fetal bovine serum (FBS) and 1% penicillin-streptomycin. WM115 and WM2664 were cultured in Minimum Essential Medium (MEM-*α*) (Invitrogen) supplemented with 1% penicillin-streptomycin (Gibco, NY, USA) and 10% FBS. NZM9, NZM40, NZM12, NZM15, NZM42 were cultured in MEM-α media supplemented with 1% penicillin-streptomycin, 5% FBS and 0.1% Insulin-transferrin-selenium (Roche)[14]. MM127, MM595 and COLO239F were grown in RPMI 1640 (ThermoFisher Scientific) Medium supplemented with 10% FBS and 1% penicillin-streptomycin. All cells were grown under standard cell culture conditions (5% CO2, 21% O2, 37°C, humidified atmosphere), except WM115, which were cultured at 35°C.

### FACs analysis

Fixable viability stain 450 (FVS450, BD horizon, catalog#: 562247, clone: 29E.2A3) was used to stain dead cells in order to selectively analyse live cells. The FVS450 stain has a fluorescence emission maximum at 450 nm. The PE anti-human CD274/PD-L1 (Biolegend, catalog# 329706) antibody and the isotype control antibody (PE Mouse IgG2b, Biolegend, catalog#:400314) has a maximum excitation at 575 nm. No overlap in fluorescence emission was detected between the FVS450 and the anti-PDL1 fluorophore or isotype control antibodies. The PD-L1 expression levels of melanoma cell lines were determined using BD FACS CantoII. All analyses were performed using the Kaluza (Beckman Coulter, version 2.0) software. Approximately 10,000 events/cells were measured for each sample. Gating strategy was used to exclude dead cells and doublet cells. The median fluorescence intensity (MFI) for the isotype control and anti-PDL1 was obtained. The MFI for PD-L1 staining was normalised for background absorbance by subtracting out the isotype fluorescence value. For IFN-γ induction of PD-L1, between 50,000 to 100,000 cells were seeded in a single well of a 24-well plate overnight with 1 mL of media. For each sample, around 10 wells were seeded in order to obtain a total amount of between 500,000 to 1 million cells. The following day, the media was removed and fresh media with IFN-γ (final concentration of 100 ng/mL, prospec, catalog#:CYT-206) was added to the cells. After 1 day of IFN-γ induction, flow cytometry was used to assess PD-L1 expression.

### RNA extraction and reverse transcription

Total RNA was isolated from melanoma cell lines using the RNeasy Mini Kit (Qiagen, catalog#:74106) following the protocol manual. This involved cell lysis, homogenisation of the lysate using the QIAshredder (Qiagen, catalog#:79656), and using a spin column to selectively purify RNA. DNase (RNase-Free DNase Set, Qiagen, catalog#:79254) was used to degrade DNA during the extraction as outlined in the RNeasy Mini Handbook. Quality control was first performed on the Nanodrop 2000 Spectrophotometer (Thermo Fisher Scientific) to assess the RNA purity using the ratio of absorbance at 260 to 280 nm higher than 1.8. The RNA integrity was assessed using the Agilent Bioanalyzer 2100 with the RNA integrity number (RIN) higher than nine. Reverse transcription from RNA to complementary DNA (cDNA) was performed using the High-Capacity cDNA Reverse Transcription Kit (Applied Biosystems, catalog#:4368814).

### Additional cell line cohorts and TCGA data for melanoma

Melanoma cell line gene expression data was obtained from external sources to validate our results. The getGEO function from the GEOquery package was used to download four gene expression datasets with a total of 175 melanoma cell lines. The datasets included: GSE7127 (n = 63) [15], GSE4843 (n = 45) [16], GSE61544 (n = 13) [17], GSE80829 (n = 54) [18]. Normalisation was performed where the unprocessed microarray data or raw count RNA-seq matrix was available. For GSE7127 (Affymetrix U133 Plus 2 microarray platform), the data was normalised using Robust Multichip Average (RMA, from the affy package). For GSE61544, the raw count data was downloaded and normalised using TMM (edgeR package)[19]. For GSE4843 the normalized data (MAS5.0 normalization) and GSE80829 (FPKM using conditional quantile normalization) data was downloaded used for analysis. RNA-seq data (FPKM) of melanoma cell lines prior to and following acquired resistance upon treatment with MAPK inhibitors as well as patient tumors that were on MAPKi treatment (4 weeks) was obtained from GSE75313 [6]. Patient tumor RNA-seq data containing baseline and after tumor progression, was obtained from GSE65186 (FPKM) (n = 70) [20]. RNA-seq data (FPKM) of baseline tumors from patients treated with anti-PD1 therapy was obtained from GSE78220 (pre-treatment tumors, n = 27) [21]. RNA-seq data (FPKM) of pre-treatment (n = 51) and on-treatment (n = 58) samples were acquired from GSE91061 [22].

### TCGA SKCM data download

The RNA-seq count matrix was downloaded from the harmonized TCGA SKCM dataset (GRCh38). This was done using the RTCGAbiolinks package in R which downloads the data from the Genomic Data Commons (GDC) database [23]. The GDCquery function was used with the parameters: project = “TCGA-SKCM”, data.category = “Transcriptome Profiling”, data.type = “Gene Expression Quantification”, workflow.type = “HTSeq - Counts”). The TMM method was used to normalise the count matrix.

### Generation and processing of Transcriptome data

RNA-seq processing for PD-L1_IND_ and PD-L1_CON_ melanoma cell lines was performed with poly-A-tail selection, paired-end reads, read length of 2x 100 bps and a total of 40 million reads. Adaptor trimming was done using cleanadaptors from the DMAP package [24,25]. The RNA-Seq design and analysis that was used in this work has been described in detail previously [26]. Pseudo-alignment methods such as Kallisto and Salmon were found to outperform other alignment methods when measuring lncRNA expression abundance [27]. Therefore, we mapped our reads to the hg38 reference genome using Kallisto (version 0.44.0) [28]. Each sample was run with 100 bootstraps and with the --bias argument to correct for potential sequence-based bias. Annotations were acquired from GENECODE (Release 28 GRCh38.p12) which entailed nucleotide sequences of all transcripts (protein-coding and lncRNA transcripts) on the reference chromosomes. Tximport was used to import the kallisto gene-level counts data into R [29].

### Statistical analysis

Following filtering our genes with low counts (lower than 1 in at least seven samples), the TMM method was used to normalise for sequencing depth and RNA composition bias using the edgeR package [19]. Differential expression analysis was performed using edgeR quasi-likelihood method [30]. A False Discovery Rate (FDR) adjusted *p* value threshold of 0.05 was used to call significant genes. Gene Set Enrichment Analysis (GSEA) was performed using gene sets available in the Broad Institute Molecular Signatures Database (MsigDB) which included H1 (hallmarK), C2 (curated) and C5 (gene ontology) [31]. CAMERA test was performed as available from the edgeR package and a FDR adjusted 5 × 10^−5^ *p* value was used as the statistically significant threshold [32]. For generating a gene-set score for each sample, single sample GSEA (ssGSEA) [33] was used from the GSVA package [34]. The IFN score, TNF score, differentiation score and oxidative phosphorylation score were obtained from MSigDB from the “MOSERLE_IFNA_RESPONSE”, “PHONG_TNF_TARGETS_UP”, “GO_MEL-ANOCYTE_DIFFERENTIATION” and “KEGG_OXIDATIVE_PHOSPHORYLATION” gene sets, respectively. The viral mimicry score was self-curated from published articles [35,36] and included *DDX58, DDX41, IFIH1, OASL, IRF7, IRF1, ISG15, MAVS, IFI27, IFI44, IFI44L* and *IFI16*. To infer cytotoxic immune activity from gene expression data, the CYT-score was calculated by finding the geometric mean from *GZMA* (granzyme A) and *PRF1* (perforin) expression values [37]. The absolute abundance of eight immune and two stromal cell populations were estimated from normalised RNA-seq data using MCPcounter [38]. R codes to calculate the moving average and generate the figures were acquired from Riesenberg and colleagues [39].

## 3. Results

### PD-L1_CON_ melanoma cell lines (high PD-L1 group) have a distinct gene expression profile compared to PD-L1_IND_ melanoma cell lines (low PD-L1 group)

Melanomas can be categorized into four subgroups based on high or low levels of tumor-associated PD-L1 protein expression, and the presence or absence of TILs in melanoma tissue. In this study, we defined these subgroups as TIL+/PDL1+ (group 1), TIL- /PDL1-(group 2), TIL+/PDL1-(group 3), and TIL-/PDL1+ (group 4) (figure 1A). However, factors leading to PD-L1 expression in melanoma tumor tissues are complicated by the fact that cytokines secreted from TILs frequently induce PD-L1 expression in tumor cells, which masks any cytokine-independent, tumor cell-intrinsic PD-L1 expression (i.e. PD-L1 constitutive expression, or PD-L1_CON_). Therefore, we used melanoma cell lines to begin our investigation, as they lack TILs, and high levels of PD-L1 expression in these cell lines consequently represent tumour cell-intrinsic mechanisms. In this study, we inferred that the PD-L1_CON_ and PD-L1_IND_ melanoma cell lines most closely represent melanoma subgroups, TIL-/PDL1+ (group 4), and TIL-/PDL1- (group 2), respectively (figure 1A). Assignment of a panel of melanoma cell lines to the high and low PD-L1 expression groups was accomplished by characterizing cell surface PD-L1 protein levels and mRNA expression levels, which were assessed using flow cytometry and RNA-Seq (Supplementary Figure 1A-B). The *CD274* (PD-L1) mRNA levels (as quantified by RNA-Seq) strongly correlated with the PD-L1 surface protein expression levels (Pearson R=0.92, *p* value =1.4e10-7 Supplementary Figure 1C) in the analysed 17 cell lines (seven PD-L1_CON_ cell lines, and ten PD-L1_IND_ cell lines). Given that, occasionally melanoma cell lines and patient tumors exhibit dysfunctional IFN signalling due to genetic defects [40-44], each of the ten low PD-L1_IND_ cell lines in our panel was additionally confirmed as being able to respond to IFNγ induction, which was verified by PD-L1 upregulation upon IFNγ treatment (Supplementary Figure 1D).

**Figure 1:**
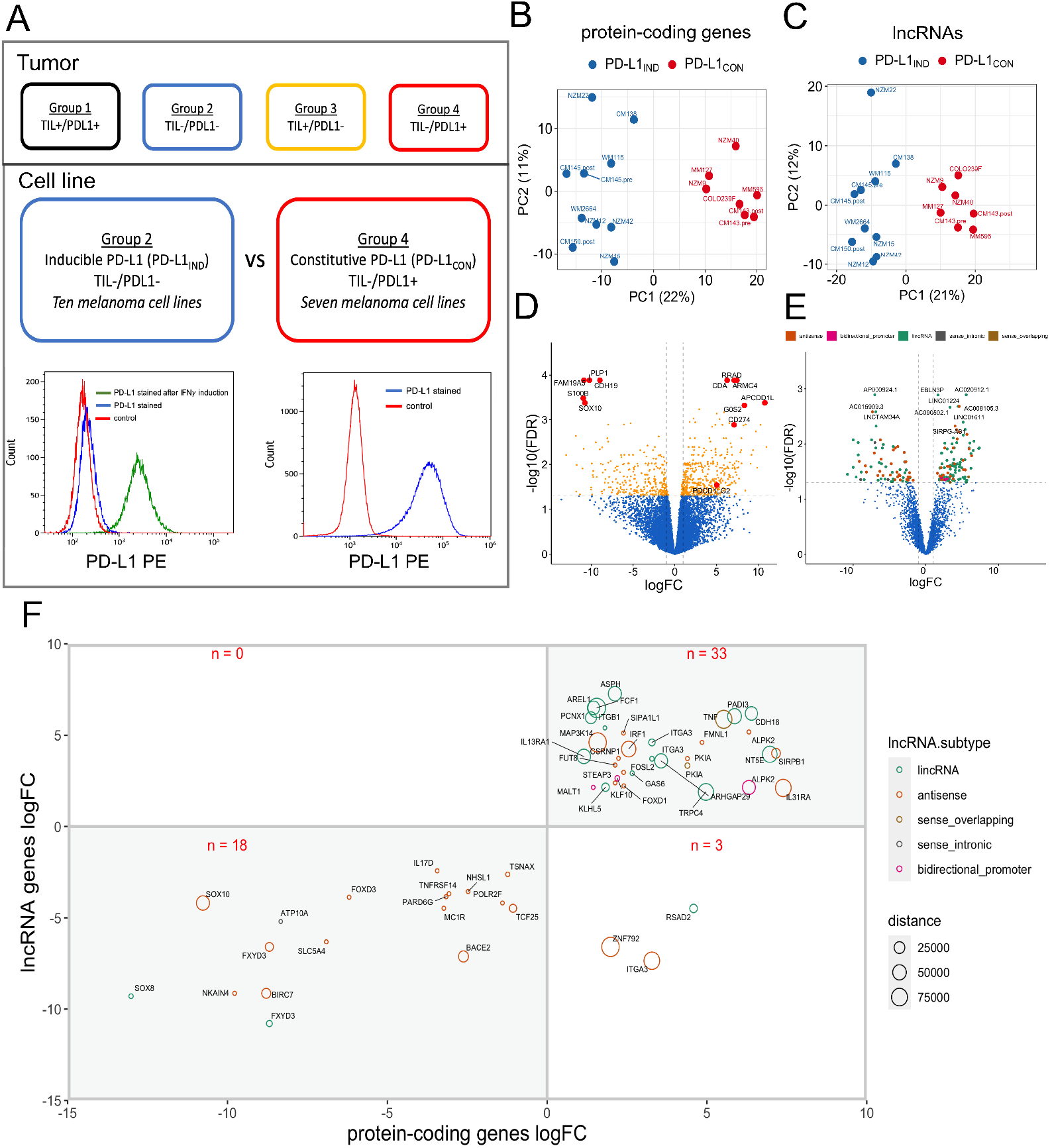
The PD-L1_CON_ group harbours distinct global mRNA and long non-coding RNA expression profiles compared to the PD-L1_IND_ group. (A) A schematic diagram showing the four subgroups of melanoma tumors based on the levels of tumor infiltrative lymphocytes and PD-L1 protein expression. Out of these four groups, we assessed the TIL negative groups by using melanoma cell lines and thereby compared the transcriptome between the low (group2: TIL-/PDL1- or PD-L1_IND_) and high PD-L1 expression group (group4: TIL-/PDL1+ or PD-L1_CON_). Representative FACs figure of the PD-L1 expression levels in each of the two groups are shown below. The FACs figure shows isotype control (red), PD-L1 stained (blue) and PD-L1 stained following 24 hour IFNγ induction (green). The x-axis represents PD-L1 expression levels (log_2_) and y-axis represents cell counts. (B-C) Principal component analysis (PCA). Shows PC1 (x-axis) and PC2 (y-axis) for the PD-L1_IND_ samples and the PD-L1_CON_ samples using the top 500 protein-coding genes (B) or lncRNAs (C) with the highest variance. (D-E) Volcano plot showing the differential expressed protein-coding genes and lncRNAs. The top 10 differentially expressed gene names are shown and moreover the *CD274* (PD-L1) and *PDCD1LG2* genes (PD-L2) are shown. The x-axis represents log_2_ fold change and the y-axis represents the -log_10_ FDR adjusted *p* value. (F) The significantly differentially expressed mRNA-lncRNA gene pairs that are located within 100 kilobases to one another are shown. The x-axis and the y-axis show the protein-coding gene and lncRNA expression fold change (log_2_), respectively. Only the protein-coding gene names are shown. The distance between the mRNA-lncRNA gene pair are represented by the size of the circle and the lncRNA subtype is shown by the colors.

Principal component analysis of the top 500 protein coding (Figure 1B) and long non-coding RNA (Figure 1C) genes with the highest variance segregated the cell lines into two distinct clusters. Unsupervised hierarchical clustering showed the same result with a clear segregation of the two groups (Supplementary Figure 2A-B), which suggests there is a distinct transcriptional state between PD-L1_CON_ and PD-L1_IND_ melanoma subgroups. Differential expression analysis identified 462 upregulated and 298 downregulated protein coding genes in the PD-L1_CON_ group, compared to the PD-L1_IND_ group (FDR adjusted *p* value < 0.05). As expected, *CD274* expression was significantly increased in the PD-L1_CON_ group (FDR adjusted *p* value = 0.001, log_2_ fold-change = 7.1, Figure 1D and Supplementary Figure 2C). *PDCD1LG2* (which encodes the PD-L2 protein, and is also located in the same chromosome location as *CD274*, i.e. 9p24.1) was also significantly upregulated in the PD-L1_CON_ group (FDR adjusted *p* value = 0.03, log2FC = 5.0, Figure 1D, Supplementary Figure 2D). For the long non-coding RNA (lncRNA) analysis, we identified 106 upregulated and 71 downregulated lncRNA in PD-L1_CON_ lines compared to the PD-L1_IND_ lines (Figure 1E). The upregulated lncRNAs included 75 lincRNAs, 26 antisense, 3 bidirectional promoter and 2 sense overlapping lncRNAs, while the downregulated lncRNAs included 37 lincRNAs, 31 antisense and 3 sense intronic lncRNAs. Next, given that lncRNAs can regulate gene expression via cis mechanisms, we assessed all differentially expressed lncRNAs with loci mapped adjacent to (within 100 kb) a differentially expressed protein-coding gene, which identified 54 mRNA-lncRNA pairs. 51 out of the 54 (94.4%) mRNA-lncRNA pairs showed expression changes in the same direction (either up or down), with 33 pairs being upregulated (top-right quadrant, Figure 1F), while 18 pairs showing downregulation (bottom-left quadrant, Figure 1G) in the PD-L1_CON_ lines compared to the PD-L1_IND_ group. These results suggest coordinated expression changes for the mRNA and lncRNA expression. Overall, PD-L1_CON_ cell lines had a distinct expression profile for protein-coding genes and lncRNAs, and moreover the vast majority of the differentially expressed mRNAs and lncRNAs located in genomic proximity to each other were altered in the same direction.

### PD-L1_CON_ cell lines exhibit a distinct transcriptome that represents a state of dedifferentiation, enhanced IFN and TNF signalling pathways and reduced oxidative phosphorylation

To determine which biological processes are altered in the PD-L1_CON_ cell lines compared to PD-L1_IND_ cell lines, we performed gene-set enrichment analysis using CAMERA [32] and C2, C5 and H gene set collections from the Molecular Signature Database (MSigDB) [31]. Thirty seven gene sets were found to be significantly altered (FDR adjusted *p* value threshold of 5 × 10^−5^). Genes involved in interferon-alpha (IFN-α) and tumor necrosis factor (TNF) signalling were among the most significantly upregulated processes in the PD-L1_CON_ group [(FDR adjusted *p* value = 3.0 × 10^−10^(MOSERLE_IFNA_RESPONSE) and 1.1× 10^−9^ (PHONG_TNF_TARGETS_UP), Figure 2A and Supplementary Figure 3A]. Given that the PD-L1_CON_ cell lines do not harbor immune cells, this suggests that the IFN and TNF signalling could be activated via the expression of dsRNA derived from endogenous retroviral elements (ERVs). Expression of ERVs can trigger dsRNA sensors and a downstream signalling cascade of interferon response genes (also referred to as a viral mimicry response) [35,36]. Genes involved as dsRNA sensors, as well as the viral mimicry response genes, were also highly upregulated in PD-L1_CON_ group (FDR adjusted *p* value = 7.5 × 10^−4^ and 2.1 × 10^−4^, respectively)(Supplementary Figure 3A). These findings suggest that PD-L1_CON_ samples may have enhanced activation of viral mimicry pathways. This is also supported in our previous findings, in that ERV genomic regions were hypomethylated in PD-L1_CON_ melanoma cells [7], and were a characteristic feature associated with PD-L1 expression. The downregulated genes in the PD-L1_CON_samples were significantly enriched for genes involved in melanocyte differentiation (FDR adjusted *p* value = 2.7 × 10^−6^, Figure 2A and Supplementary Figure 3A), suggestive of a dedifferentiated state. Recently, Tsoi and colleagues found that melanomas can exhibit transcriptomic states that are coupled to a differentiation trajectory, which consists of four progressive steps [18], corresponding to 1) undifferentiated, 2) neural crest-like, 3) transitory and 4) melanocytic. Unsupervised hierarchical clustering of the cell lines based on these melanoma differentiation genes confirmed that the PD-L1_CON_ cell lines were enriched for genes characteristic of the undifferentiated state (Figure 2B). The PD-L1_IND_ cell lines were further clustered into two subgroups from their transcriptomic profiles, with one group corresponding to a neural crest-like gene signature, and the other group to a melanocytic state. Moreover, there was a downregulation of genes involved in oxidative phosphorylation in the PD-L1_CON_ group suggesting metabolic reprogramming had occurred (FDR adjusted *p* value = 7.2 × 10^−8^ (KEGG_OXIDATIVE_PHOSPHORYLATION)) (Figure 2B and Supplementary Figure 3A). Given that dedifferentiation is closely associated with enhanced invasiveness we investigated whether the PD-L1_CON_ samples were enriched for an invasive gene expression profile. Using two independent invasive gene expression gene sets [45,46], we found PD-L1_CON_ cell lines had higher levels of expression of genes involved in melanoma invasion (Supplementary Figure 4A-B) although three PD-L1_IND_ cell lines (NZM22, WM115, CM145.pre) had weak expression levels of invasive genes, whereas CM138 clustered more closely to the PD-L1_CON_ cell lines.

**Figure 2.**
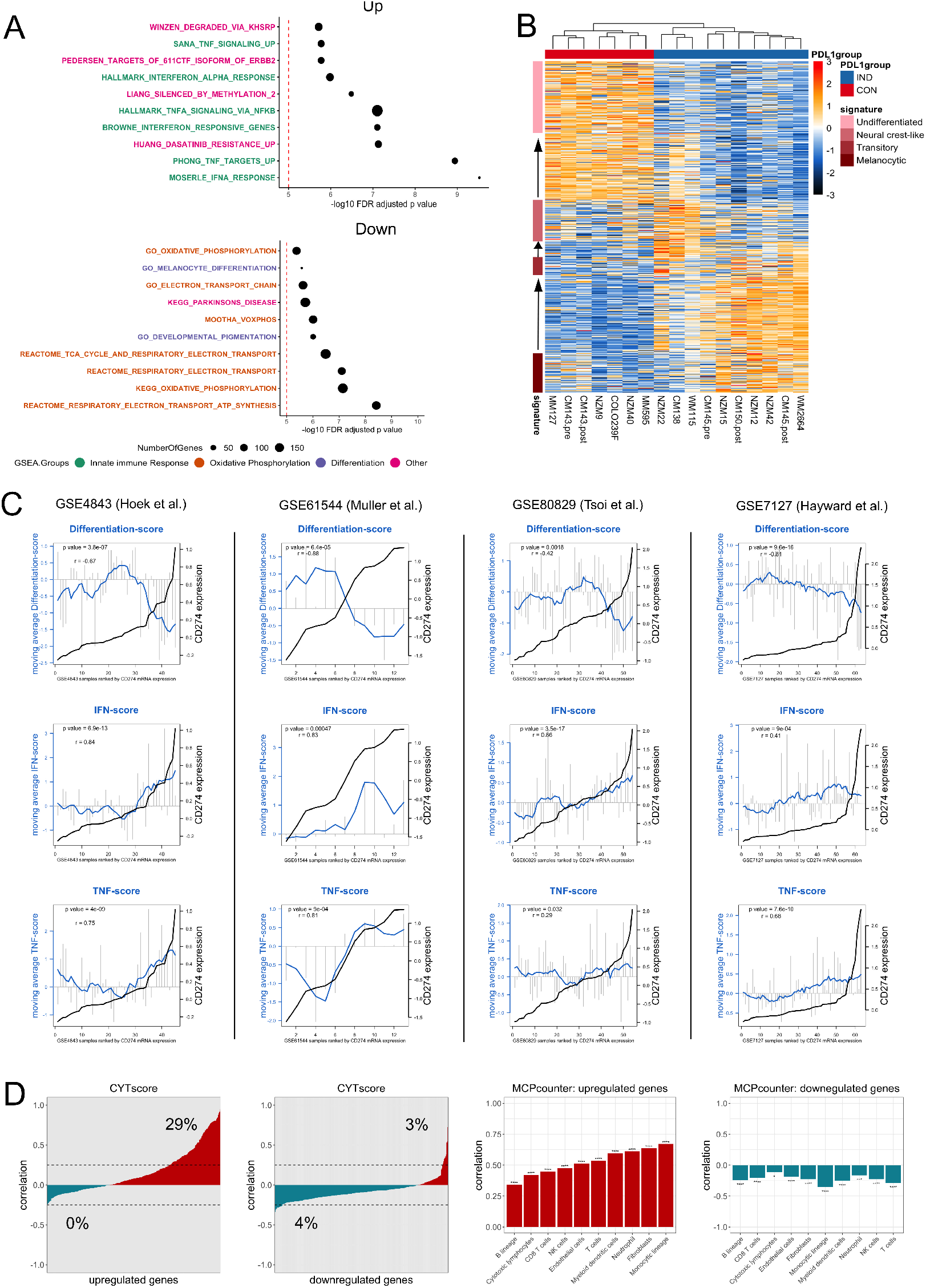
PD-L1_CON_ samples have a reprogrammed transcriptome inclusive of dedifferentiation, reduced oxidative phosphorylation and an enhanced IFN signalling pathway. (A) The top 20 gene sets that was found to be differentially enriched using the CAMERA test are shown [32]. The 5 × 10- 5 FDR adjusted *p* value significance threshold is shown by the dotted red line. Gene sets are categorised into those involved in the innate immune response (or cytokine signalling), melanocyte differentiation, oxidative phosphorylation and others. (B) Heatmap showing the unsupervised clustering of the PD-L1_CON_ and PD-L1_IND_ melanoma cell lines according to the differentiation signature genes from Tsoi and colleagues [18]. (C) Correlation of *CD274* expression with the IFN score, TNF score and differentiation score in the four external melanoma cell line datasets. The samples (x-axis) are ranked according to *CD274* expression (log_2_ transformed and corresponds to the right side y-axis) and the *CD274* expression level is shown by the black line. Grey vertical bars indicate the respective scores and corresponds to the left side y-axis. The moving average is represented by the blue line. The Pearson correlation between the moving average and *CD274* expression is shown. (D, first and second figure) The y-axis represents the correlation value (Pearson) of the CYT score with either the upregulated (D, first figure) or downregulated protein-coding genes (D, second figure) in the PD-L1_CON_ samples. The x-axis represents all the up (462 genes) and downregulated (298 genes) protein-coding genes. The x-axis is ranked according to lowest to highest correlation value (Pearson). (D, third and fourth figure) The y-axis represents the correlation value (Pearson) of the up (D, third figure) and down PD-L1_CON_ scores (D, fourth figure) (calculated with ssGSEA using the differentially expressed protein-coding genes) with the estimated absolute abundance of various immune subtypes (as measured using the MCPcounter bioinformatic tool).

### Validation of upregulated IFN and TNF pathways, and downregulated differentiation and oxidative phosphorylation pathways in association with constitutive *CD274* expression in melanoma cell lines

To validate the gene expression signature of an upregulated IFN and TNF pathway, and downregulation of differentiation and oxidative phosphorylation genes in PD-L1_CON_ melanoma cells, we utilised four external gene expression datasets containing a total of 175 melanoma cell lines. A score was generated for the 462 upregulated and 298 downregulated genes in the PD-L1_CON_ samples (referred to as the up and down PD-L1_CON_ score respectively) as well as the genes involved in IFN signalling, TNF signalling, differentiation and oxidative phosphorylation using single sample gene set enrichment analysis (ssGSEA) [33,34]. In all four external gene expression datasets, *CD274* expression was significantly positively correlated with IFN and TNF scores and negatively correlated with the differentiation score (Figure 2C). However, the oxidative phosphorylation score did not show consistent correlation with *CD274* expression in the external gene expression datasets (Supplementary Figure 3B).

The upregulated IFN and TNF signalling in the PD-L1_CON_ samples which are occurring cell intrinsically are similar to tumors where immune cell infiltrates secrete cytokines to extrinsically induce IFN signalling and PD-L1 expression. To further investigate the upregulated IFN and TNF signalling in the PD-L1_CON_ samples, we asked whether the PD-L1_CON_ samples share an expression signature with those tumors with an elevated immune infiltration. To this end, the TCGA SKCM dataset which contains RNA-seq data for 472 tumours was analysed. A large proportion of significantly upregulated transcripts (135 out of 462 or 29%) in the PD-L1_CON_ samples were positively correlated (Pearson correlation value of higher than 0.25) with the CYTscore (Figure 2D), which is a well-established index that reflects cytolytic immune activity [37]. Moreover, the upregulated and downregulated PD-L1_CON_ genes were positively and negatively correlated, respectively, with the absolute abundance of a large range of immune subtypes as estimated by MCPcounter [38] (Figure 2D), demonstrating that PD-L1_CON_ melanoma cell lines have an innately active cytokine signalling pathway similar to melanoma tumours where immune infiltrates stimulate this pathway.

### Validation of upregulated IFN and TNF pathways, and downregulated differentiation and oxidative phosphorylation pathways in association with constitutive *CD274* expression in melanoma tumour tissues

To validate the association of PD-L1_CON_ expression with active IFN and TNF signalling, and reduced differentiation and oxidative phosphorylation expression signatures in melanoma tumors, we used the SKCM TCGA dataset consisting of 458 melanoma tumour samples. As mentioned above, *CD274* expression in melanoma tissues is typically stimulated by immune infiltrates rather than an intrinsic mechanism, therefore we used the significantly differentially expressed genes that were found in the PD-L1_CON_ cell lines as a surrogate for PD-L1_CON_ expression. A large proportion of the upregulated protein-coding genes (n = 462) found in the PD-L1_CON_ samples were positively correlated (Pearson r > 0.25) with *CD274* expression (136 out of 462 or 29%), IFN score (136 out of 462 or 29%) and TNF score (213 out of 462 or 46%). Furthermore, the upregulated proteincoding genes were negatively correlated (lower than -0.25 Pearson correlation value) with the differentiation (137 out of 462 or 30%) and oxidative phosphorylation scores (151 out of 462 or 33%) (Figure 3A and Supplementary Figure 3C). In contrast, a large proportion of the downregulated protein-coding genes (n = 298) were positively correlated (Pearson r > -0.25) with the differentiation-score (127 out of 298 or 43%) and the oxidative phosphorylation score (108 out of 298 or 37%) (Figure 3A and Supplementary Figure 3C). The same pattern was also seen for the lncRNAs (Figure 3B). Collectively, these results support the finding that PD-L1_CON_ expression is associated with an enhanced IFN and TNF signalling, and reduced differentiation and oxidative phosphorylation.

**Figure 3.**
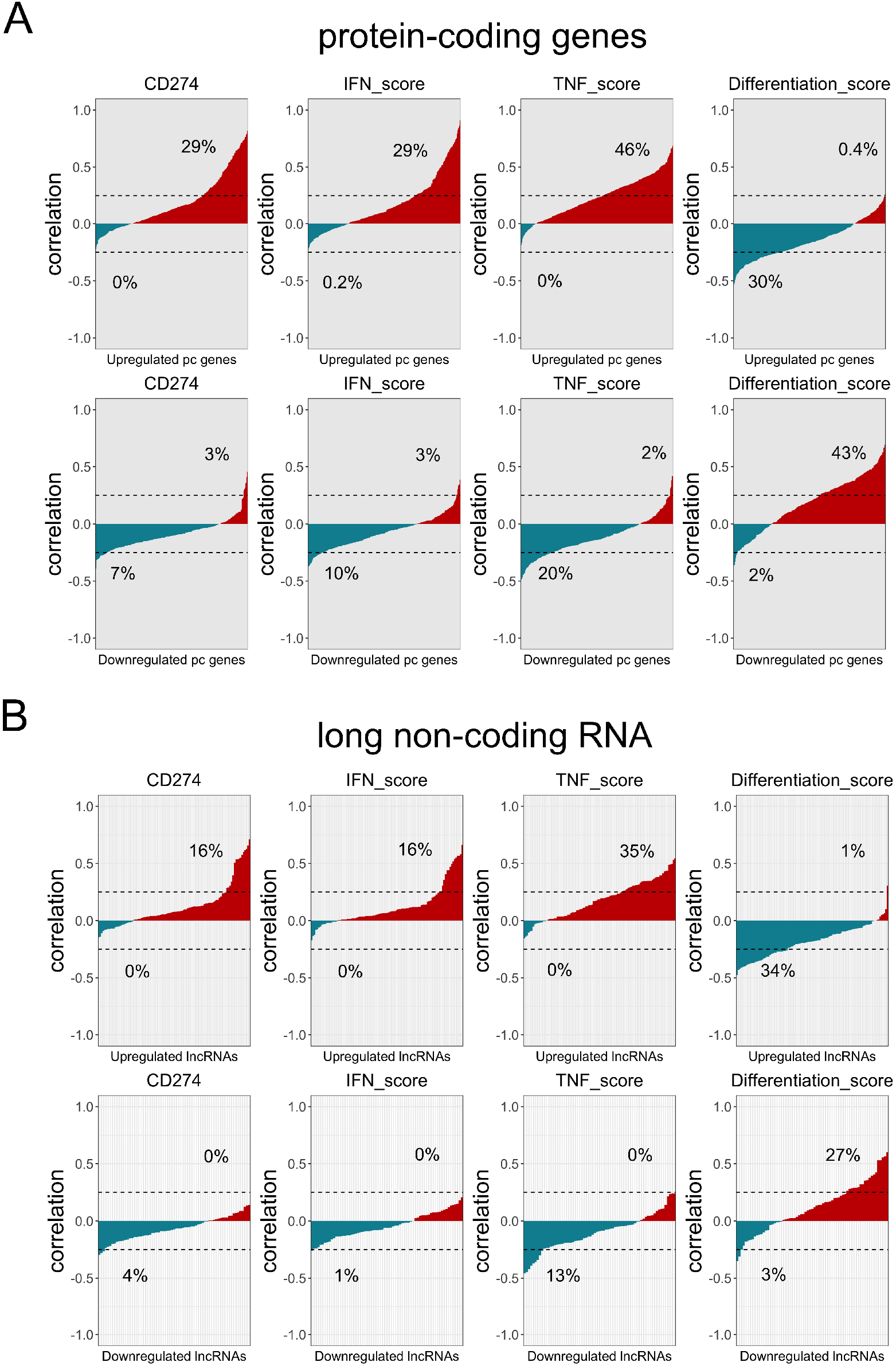
Protein-coding and lncRNA expression signatures of PD-L1_CON_ samples are associated with an increased IFN score and reduced differentiation and oxidative phosphorylation scores in the TCGA SKCM database. (A) The x-axis shows the upregulated (n = 462)(A, top row) and down-regulated (n = 298) (A, bottom row) protein-coding genes (A) in the PD-L1_CON_ samples. The y-axis shows the correlation (Pearson) of the protein-coding genes with *CD274* expression, IFN score, differentiation score and oxidative phosphorylation score. The x-axis is ranked according the lowest to highest correlation. The percentage of genes higher than 0.25 or lower than -0.25 Pearson correlation value is shown. (B) The x-axis shows the upregulated (n = 106) and downregulated (n = 71) lncRNAs in the PD-L1_CON_ samples. The y-axis shows the correlation (Pearson) of the lncRNAs with *CD274* expression, IFN score, differentiation score and oxidative phosphorylation score. The x-axis is ranked according the lowest to highest correlation. The percentage of genes higher than 0.25 or lower than -0.25 Pearson correlation value is shown.

### Lineage specific, TNF and IFN associated transcription factors are differentially expressed in the PD-L1_CON_ samples

Key transcription factors (TFs) can play a role in modulating the transcriptomic program. Therefore, we investigated what type of TFs are altered in expression in the PD-L1_CON_ cell lines. To this end, we analysed 1,107 TF mRNAs with known DNA binding motifs [47] and identified 25 significantly upregulated TF mRNAs (out of 462 significantly upregulated genes) and 19 significantly downregulated TF mRNAs (out of 298 significantly downregulated genes) in the PD-L1_CON_ cell lines (Figure 4A). To determine whether the expression patterns of these TF mRNAs could be generalised to melanoma, we analysed the four external gene expression datasets and the TCGA SKCM RNA-seq dataset in the TF context. The 25 upregulated TF and 19 downregulated TFs were predominantly positively and negatively correlated with *CD274* expression in all five datasets (Figure 4B). Furthermore, we identified *POU2F2, IRF1* and *FOSL2* as the most highly correlated TFs with *CD274* expression in melanoma across all five datasets (Figure 4C). IRF1 and FOSL2 have known roles in upregulating PD-L1 expression [9,48,49]. In contrast, *SOX10, RXRG* and *SOX5* had the lowest correlation with *CD274* expression amongst all TFs investigated (Figure 4C). SOX10 and MITF are key transcription factors that regulate the development of melanocytes. *SOX10* expression showed a near complete loss (log2FC = -10.7, FDR adjusted *p* value = 0.0003) and *MITF* expression was significantly reduced in expression (log2FC= -3.8, FDR adjusted *p* value = 0.04) in the PD-L1_CON_ cell lines (Supplementary Figure 5E-F). The upregulation of these TF expression further supports our observation that the PD-L1_CON_ melanoma subgroup is associated with substantial transcriptomic reprogramming.

**Figure 4.**
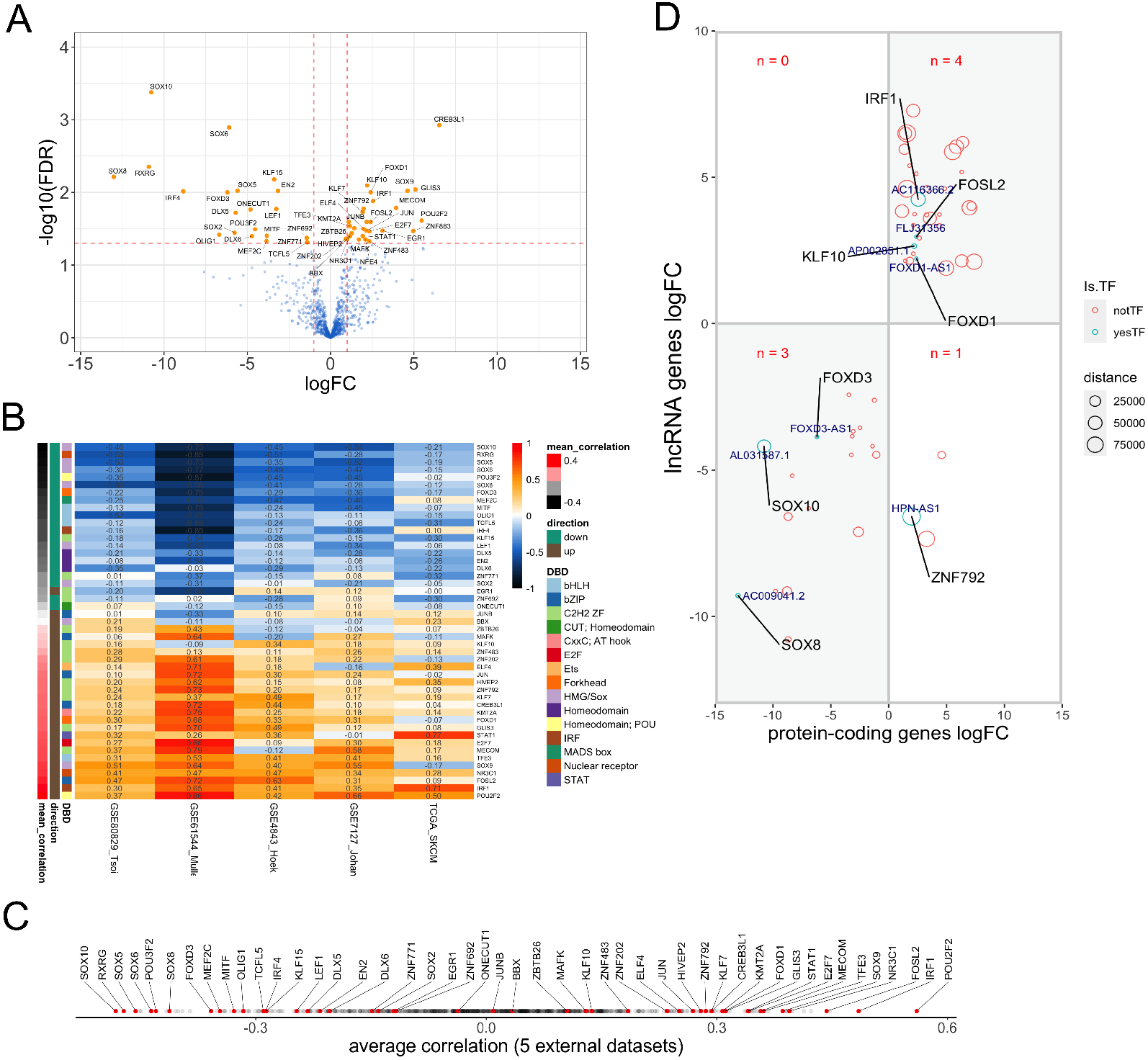
Correlation of lineage-specific and immunity-associated transcription factor mRNAs with PD-L1 expression in melanoma and association with long non-coding RNAs. (A) Volcano plot showing all the differentially expressed TFs in the PD-L1_CON_ group. The 25 and 19 significantly upregulated and downregulated TF mRNAs in the PD-L1_CON_ samples are shown respectively. The x-axis represents log_2_ fold change and the y-axis represents -log_10_ FDR adjusted *p* value. The red dashed line represent the FDR adjusted 0.05 *p* value (y-axis) and the 1 log_2_ fold change (x-axis). (B) Heatmap showing the correlation value (Pearsons) of the 25 up and 19 down regulated TFs with *CD274* expression. Rows represent the TFs and are ordered according to the lowest to highest mean correlation with *CD274* expression across the four melanoma cell line datasets and the TCGA SKCM RNA-seq dataset. (C) Correlation was calculated between *CD274* expression and all TF (n = 1,107) across the five gene expression datasets and the TF were ordered from highest to lowest mean average correlation across the five datasets. The direction of the 25 upregulated and 19 downregulated TFs are shown by the direction sidebar. (D) The significantly differentially expressed TFs to lncRNA pairs that are located within 100 kilobases to each other are shown. The x-axis and the y-axis show the TF mRNA and lncRNA expression fold-change (log_2_), respectively. The distance between the TF mRNA and lncRNA gene are shown by the size of the circle.

Additionally, we identified all differentially expressed lncRNAs whose loci were located in close proximity/ adjacent to differentially expressed TF mRNAs (within 100kb of each other). Correspondingly, eight TF mRNA and lncRNA differentially expressed pairs were found (Figure 4D). Out of these eight TF’s, four were identified to be closely associated with *CD274* expression in the five external datasets. This included *AL031587*.*1* antisense lncRNA near *SOX10* (distance of 44,743 bp), the *AC116366*.*2* intergenic lncRNA near *IRF1* (distance of 54,895 bp) and FLJ31356 antisense lncRNA near *FOSL2* (distance of 0 bp, as they overlap)(Figure 4D).

### The PD-L1_CON_ expression signature is associated with transriptomic reprogramming, and correlates with MAPK inhibitor resistance in melanoma cell lines

Key transcriptional changes that were shown to drive acquired resistance include increased expression of *cMET*, reduced expression of *LEF1*, and an enrichment of YAP1 signature genes [20]. PD-L1_CON_ cell lines had increased expression of *cMET* (log2FC = 2.6, FDR adjusted *p* value = 0.07), reduced expression of *LEF1* (log2FC = -3.2, FDR adjusted *p* value = 0.017) and enrichment of YAP1 signature genes (t.test *p* value = 0.012) (Supplementary Figure 5B-D). Other gene expression changes driving drug resistance include reduced levels of *MITF* and *SOX10* which were both downregulated in the PD-L1_CON_ cells. Moreover, consistent with increased expression of Receptor Tyrosine Kinases (RTK), which are associated with drug resistance, we found a large number of RTK genes were highly expressed in the PD-L1_CON_ cell lines including *EGFR* (log2FC= 4.2, FDR adjusted *p* value = 0.054), *PDGFRB* (log2FC= 3.6, FDR adjusted *p* value = 0.03) and *AXL* (log2FC= 2.5, FDR adjusted *p* value = 0.34) (Supplementary Figure 5A and 5G-I). These data support the notion that PD-L1_CON_ cell lines have a transcriptional profile corresponding to a drug resistant phenotype.

*CD274* expression was found to be cell-intrinsically increased upon development of resistance to MAPK inhibitors, when accompanied by transcriptomic reprogramming, but not mediated by *BRAF* mutations (splicing or amplification) reactivating the MAPK pathway [6]. We utilised this dataset to assess our PD-L1_CON_ gene expression signature in the context of development of resistance. In cell lines where resistance co-occurred with transcriptomic reprogramming, there was an increase of *CD274* expression and as well as IFN, TNF and “up” PD-L1_CON_ scores (Figure 5B), and a corresponding reduction in differentiation, oxidative phosphorylation, MPAS and “down” PD-L1_CON_ scores. Moreover, we found that TFs associated with PD-L1_CON_ expression, and involved in IFN signalling and TNF signalling (*IRF1, JUN* and *FOSL2*), were all upregulated (Figure 5E), except for *IRF4*, which was downregulated in these cell lines. Consistently, *IRF4* was the only TF involved in IFN and TNF signalling that was also downregulated in PD-L1_CON_ cell lines (Figure 5C). In addition, TFs involved in melanocyte differentiation (including *SOX10, MITF* and *LEF1*) were downregulated in cell lines that the authors had associated with the development of drug resistance (Figure 5E). In contrast, in cell lines where resistance was mediated by reactivation of the MAPK signalling pathway (involving *BRAF* splicing or amplification), only minimal changes in the expression of *CD274* and the PD-L1_CON_ related biological processes (including IFN signalling, TNF signalling, differentiation, and oxidative phosphorylation) were observed (Figure 5A). Additionally, the expression of TFs involved in IFN or TNF signalling, or melanocyte differentiation were unaltered when the drug resistance was mediated by *BRAF* splicing or amplification mutations (figure 5D).

**Figure 5.**
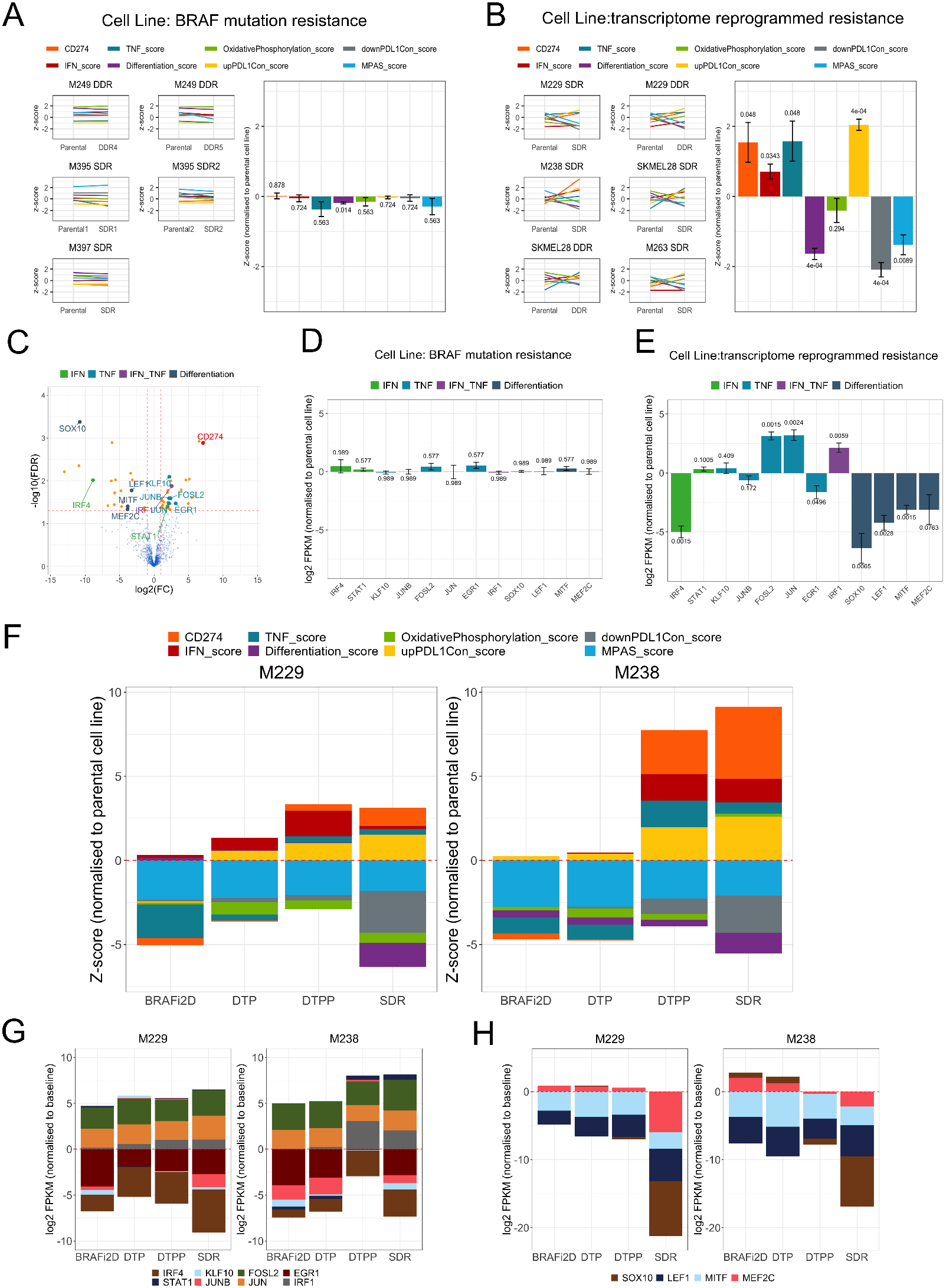
PD-L1_CON_ expression is associated with treatment resistance to MAPK inhibitors when associated with transcriptome reprogramming. (A-B) Five melanoma cell lines that acquired BRAF mutations (splicing and copy number amplification) upon development of resistance and six melanoma cell lines that did not acquire mutations but exhibited a transcriptome reprogrammed state upon development of resistance are shown. *CD274* expression and seven scores which includes the IFN score, TNF score, differentiation score, oxidative phosphorylation score, the MPAS score, and the up and down PD-L1_CON_ scores are shown for the parental (before treatment) and after BRAFi resistance (SDR = single drug resistance, DDR = double drug resistance). The bar plots show values normalised to the corresponding baseline sample. Bar plot values are averages across all samples within either the BRAF mutation resistance group or the transcriptome reprogrammed resistance group. FDR adjusted *p* values for paired t test are shown for each bar. Error bars represents standard error. (C) Volcano plot shows the TFs that were differentially expressed in the PD-L1_CON_ samples and in addition, involved in IFN signalling, TNF signalling, both IFN and TNF signalling (labelled as “IFN_TNF”) and the melanocyte differentiation pathway. The *CD274* gene is shown in red. (D-E) Bar plots shows the expression changes of TFs (involved in the IFN, TNF signalling and melanocyte differentiation) following acquired resistance with either BRAF mutations or transcriptome re-programming in melanoma cell lines. Error bars represents standard error. (F) Cumulative barplot shows the *CD274* expression and the seven scores at different stages of BRAFi resistance for the M229 and M238 melanoma cell lines. Values are normalised to the parental cell line and resistance stages include (1) two days of BRAFi treatment (BRAFi2D), (2) Drug Tolerant Persisters (DTP) where a small subpopulation of persisting cells remain, (3) Drug Tolerant Proliferative Persisters (DTPP) where proliferation was regained and (4) single drug resistance (SDR), a permanent resistant state to BRAFi. (G-H) Cumulative barplot shows the mRNA expression changes of TFs involved in the IFN signalling, TNF signalling and melanocyte differentiation at different stages of BRAFi resistance for the M229 and M238 melanoma cell lines.

### Transcriptomic changes, including PD-L1_CON_ expression, occur at defined stages during the development of drug resistance

To investigate the timing of when PD-L1 expression changes and biological processes may occur during drug treatment, we investigated RNA-Seq data derived from two melanoma cell lines (M229 and M238)[6], which had been treated with an MAPK inhibitor, and for which RNA-Seq analysis was carried out at different time points between the initial resistance, generation of drug tolerance, and finally permanent acquired resistance. The time points included after two days of BRAFi treatment (BRAFi2D); at the Drug Tolerant Persisters (DTP) stage where small subpopulation of persisting cells remained; at the Drug Tolerant Proliferative Persisters (DTPP) stage where proliferation was regained; and finally at the Single Drug Resistant (SDR) stage, which is a permanent resistant state to BRAFi (months to years of drug treatment). *CD274* expression was slightly decreased at the DTP stage however increased at the proliferative phase (DTPP) and was further increased at the SDR phase (Figure 5F). TNF, IFN and PD-L1_CON_ “up” scores were largely increased at the DTPP stage, whereas dedifferentiation and PD-L1_CON_ “down” scores were largely decreased at the SDR stage. This suggested that innate IFN and TNF signalling responses precede dedifferentiation. Further analysis revealed that *JUN* and *FOSL2*, which encode proteins that are part of the AP-1 complex, and are part of the TNF signalling pathway, were increased relatively early after two days of BRAFi treatment and were maintained until acquired resistance developed, whereas *IRF1*, which can drive PD-L1 expression, increased at the DTPP stage when *CD274* was also overexpressed (Figure 5G). Furthermore, the dedifferentiation TFs, including *MITF* and *LEF1* were downregulated early after two days of BRAFi treatment, whereas *SOX10* and *MEF2C* were reduced upon the development of acquired resistance (SDR), which was also when the differentiation score was greatly reduced. This suggested that *SOX10* and *MEF2C* downregulation are required, or are the main contributors to the dedifferentiation gene expression signature. Overall, these data support the notion that PD-L1_CON_ expression occurs at the proliferative stage of drug resistance and that this is mediated by a transcriptome reprogramming, and is accompanied by increases in TNF and IFN signalling along with the *IRF1* expression. Moreover, these expression changes further increase when resistance is stabilised, at which point the differentiation expression signature is downregulated.

### The PD-L1_CON_ expression signature is associated with transcriptomic reprogramming of melanomas following MAPK inhibitor resistance in patients

We next investigated whether the PD-L1_CON_ expression signature occurs in melanomas from patients who develop acquired resistance to MAPK inhibitor treatment. To address this, we analysed RNA-Seq data from matched pre- and post-treatment tumor biopsies, following progression on treatment [20]. We compared melanomas with *BRAF* mutational resistance mechanisms (*BRAF* splicing and amplification) to melanomas lacking known mutations that drive resistance. In melanomas where no mutational resistance mechanism was identified, we observed the same direction of up and down gene expression changes to the PD-L1_CON_ cell lines, and to the transcriptomic changes of Song and colleagues’ dataset, where resistance was mediated by transcriptomic reprogramming. This consisted of elevated levels of *CD274* expression, and elevated IFN, TNF, and the “up” PD-L1_CON_ scores (normalised to matched pre-treatment tumors from the same patient), while oxidative phosphorylation, down PD-L1_CON_, and differentiation scores were reduced (Supplementary Figure 6A). In addition, there were lower levels of the differentiation related TFs and higher levels of expression of *JUN, IRF1, JUNB* and *FOSL2* observed in melanomas with no *BRAF* mutational resistance mechanisms (normalised to patient matched pre-treatment tumors), compared to melanomas containing *BRAF* mutational resistance mechanisms (Supplementary Figure 6B). These results support the notion that the PD-L1_CON_ expression signature accompanies transcriptomic reprogramming and treatment resistance in melanomas lacking a *BRAF* mutational resistance mechanism.

## 4. Discussion

The use of targeted therapies based on MAPK pathway inhibition, or of anti-PD1 immunotherapy, has improved melanoma patient survival, although drug resistance greatly limits long-term survival for most patients with advanced melanoma. A large number of mutational and non-mutational (transcriptomic or epigenetic) resistance mechanisms have been identified in melanoma [6,20,21,50]. Moreover, these have been associated with a cell-intrinsic upregulation of the immunosuppressive PD-L1 protein [5,6]. In the present study, to explore how PD-L1 expression is associated with transcriptomic reprogramming and drug resistance, we have assessed the transcriptomic landscape of melanoma cell lines with constitutively high levels of PD-L1 expression (PD-L1_CON_). Overall, our study has shown that PD-L1_CON_ cells have many similarities to melanoma cell lines and patient tumors that exhibit intrinsically upregulated PD-L1 expression following the development of resistance to MAPK pathway inhibitors via non-mutational and/or epigenetic mechanisms. Our research reveals that this similarity is inclusive of a gene expression profile corresponding to dedifferentiation, and active cytokine (TNF and IFN) signaling pathways. Dedifferentiation is a non-mutational mechanism of drug resistance, which melanomas commonly exploit. Consistent with other studies showing that treatment of melanoma and melanocytes cells with TNF-α and IFN-γ cytokines can induce a dedifferentiated state, we found that the dedifferentiated state of PD-L1_CON_ cells is associated with constitutively active IFN and TNF signaling pathways [18,39,51,52]. Furthermore, given that the PD-L1_CON_ and PD-L1_IND_ cell line groups used in this study were classified into these groups independent of mutation status (four cell lines were *BRAF* mutant, two *NRAS* mutant cell lines, and one *BRAF*/*NRAS* wild-type), these data suggest that the presence of genetic mutations did not influence the occurrence of a PD-L1_CON_ transcriptomic expression pattern, and lend support to the notion that transcriptional or epigenetic factors can over-ride mutation-driven resistance in the absence of a targeted drug treatment.

It is important to note that almost all of the PD-L1_CON_ melanoma cell lines (six out of seven) in this study had not been previously exposed to targeted inhibitors either in the patient (*in vivo*) or *in vitro*. Nevertheless, each PD-L1_CON_ cell line contained the “resistant” gene expression signature. This suggests that stress factors, other than targeted inhibitor drugs, may have induced the resistant transcriptional profile. This could include stresses, such as hypoxia, or treatment using chemotherapeutic drugs in the patients before the cell lines were initiated. The clinical relevance of these findings is that, in some patients, PD-L1_CON_ cells may occupy a subpopulation of cells in the tumor prior to targeted inhibitor therapy, which could then result in innate resistance. Moreover, one of the PD-L1_CON_ melanoma cell lines, which was previously exposed to MAPK pathway inhibitors (i.e. CM143-post), had a related cell line from the same patient, which also exhibited PD-L1_CON_ expression prior to treatment exposure (CM143-pre), suggesting that, in this patient, a PD-L1_CON_ transcriptomic pattern may have represented a pre-existing subpopulation of cells that was present prior to treatment, and which gave rise to the drug-resistant phenotype.

## 5. Conclusions

Our study offers new insights into transcriptomic patterns associated with PD-L1_CON_ melanoma cell lines, the expression of relevant TFs, and reprogramming of melanoma cells towards a drug resistant phenotype. For instance, SOX10 and MITF are master TF regulators of melanocyte differentiation, whereby their downregulation can promote de-differentiation and confer resistance to MAPK pathway inhibitors. This then results in activation of oncogenic signaling pathways, providing alternatives to the MAPK pathway, such as pathways driven by RTKs [17,50,53-56]. Both SOX10 and MITF were significantly downregulated in the PD-L1_CON_ cells, along with a dedifferentiation gene expression profile and increased expression of RTK genes which highlights the importance of PD-L1_CON_ expression during adaptive drug resistance. Moreover, TFs that were significantly upregulated consisted of AP-1 (*JUN* and *FOSL2*) which have recently been shown to, not only mediate the TNF signalling pathways, but also to alter the transcriptional and enhancer chromatin landscape, by serving as pioneer TFs [57-59]. Transcriptomic reprogramming and differentiation often involves pioneer TFs, which facilitate and alter access to distinct regulatory elements, and consequently alter the chromatin landscape [60]. Moreover, SOX9 has also been shown to be a pioneer TF that mediates stem cell plasticity [61-63], and we found this was significantly upregulated in the PD-L1_CON_ cells.

## Supplementary Materials

The following are available online at www.mdpi.com/xxx/s1,

### Supplementary Figure legends

Supplementary Figure 1. (A-B) PD-L1 protein and mRNA expression in the PD-L1_IND_ samples (n = 10) in comparison to the PD-L1_CON_ (n = 7) melanoma cell lines. The PD-L1 protein levels are in Median Fluorescence Intensity (MFI) normalized to isotype control by subtraction. The mRNA expression levels are log_2_ transformed TMM normalized values. (C) Correlation between the PD-L1 protein and mRNA expression. (D) PD-L1 protein levels before (control group) and after IFN γ treatment for all ten PD-L1_IND_ samples. The paired t test *p* value is shown.

Supplementary Figure 2. (A-B) Hierarchical clustering of PD-L1_CON_ and PD-L1_IND_ melanoma cell lines using either all expression of protein-coding genes or lncRNA. (C) The *CD27*4 (log_2_ fold change = 7.1, FDR adjusted *p* value = 0.001) and (D) *PDCD1LG2* mRNA expression (log_2_ fold change = 5.0, FDR adjusted *p* value = 0.03) in the PD-L1_CON_ in comparison to the PD-L1_IND_ melanoma cell lines.

Supplementary Figure 3. (A) Barcode plots showing eight gene sets with differential expression changes in the PD-L1_CON_ melanoma cell lines (red or right side) compared to the PD-L1_IND_ cell lines (blue or left side). The x-axis shows the ranked list of all protein-coding genes according to log_2_ fold-change. The vertical bars on the x-axis represent the genes that are included in the genesets. The y-axis shows the enrichment score. The FDR adjusted *p* values using the CAMERA test are shown. (B) Correlation of *CD274* expression with the oxidation phosphorylation score in the four external melanoma cell line datasets. The samples (x-axis) are ranked according to *CD274* expression (log_2_ transformed and corresponds to the right side y-axis) and the *CD274* expression level is shown by the black line. Grey vertical bars indicate the oxidative phosphorylation scores and corresponds to the left side y-axis. The moving average of the oxidative phosphorylation score is represented by the blue line. The Pearson correlation between the moving average and *CD274* expression is shown. (C) The x-axis shows the significantly upregulated (n = 462), downregulated (n = 298) protein-coding genes and the upregulated (n = 106) and downregulated (n = 71) lncRNAs in the PD-L1_CON_ samples compared to the PD-L1_IND_ samples. The y-axis shows the correlation (Pearson) of the differentially expressed genes with the oxidative phosphorylation score. The x-axis is ranked according the lowest to highest correlation. The percentage value of genes higher than 0.25 or lower than -0.25 Pearson correlation value out of all genes in the respective gene set are shown.

Supplementary Figure 4. Hierarchical clustering of the PD-L1_CON_ and PD-L1_IND_ melanoma cell lines using melanoma invasive expression signature genes. The gene signatures were derived from two independent studies: (A) Widmer et al. [46] and (B) Jeff et al. [45].

Supplementary Figure 5. (A) Hierarchical clustering of the PD-L1_CON_ and the PDL1_IND_ samples using receptor tyrosine kinase genes. Receptor tyrosine kinase genes that were differentially expressed in the PD-L1_CON_ cell lines with an FDR adjusted *p* value < 0.05 are indicated by * whereas between 0.05 and 0.1 are indicated by +. (B-I) mRNA expression levels of in the PD-L1_CON_ samples compared to the PD-L1_IND_ samples of various genes that have been found to confer resistance to BRAFi in BRAF mutant melanomas. FDR adjusted *p* values are shown.

Supplementary Figure 6. (A) Expression levels of *CD274* and the seven scores related to PD-L1_CON_ expression in the tumors that gained BRAF mutations upon acquired resistance (labelled “BRAF_mutation”) in comparison to tumors that did not gain any known mutational resistance (labelled “No_mutation”). (B) Expression levels of 12 TFs in the tumors that gained BRAF mutations upon acquired resistance (labelled “BRAF_mutation”) in comparison to tumors that did not gain any known mutational resistance (labelled “No_mutation”). All values were normalised to the baseline (pre-treatment) tumor from the same patient. The t-test *p* value is shown.

## Author Contributions

Conceptualization, A.C., E.R., P.H. and M.E.; methodology, A.A.; software, G.G., P.S..; formal analysis, A.A..; resources, P.H., M.E..; data curation, A.A., G.G., M.P.; writing— original draft preparation, A.A.; writing—review and editing, A.A., E.R., G.G., P.S., M.P., P.H., A.C., M.E.; funding acquisition, A.C., M.E. All authors have read and agreed to the published version of the manuscript.

## Data Availability Statement

DNA methylation and transcriptomic data for PD-L1_CON_ and PDL1_IND_ cell lines are available at Database: NCBI GEO, accession number <u>GSE107622</u>.

## Acknowledgments

The authors thank Drs Sarah Diermeier and Nicole Cloonan for comments and suggestions, and Professor Glen Boyle for providing melanoma cell lines.

