## Supplementary figures and images for "Transcriptional reprogramming and constitutive PD-L1 expression in melanoma are associated with dedifferentiation and activation of interferon and tumor necrosis factor signaling pathways"

### Supplementary Figure 1

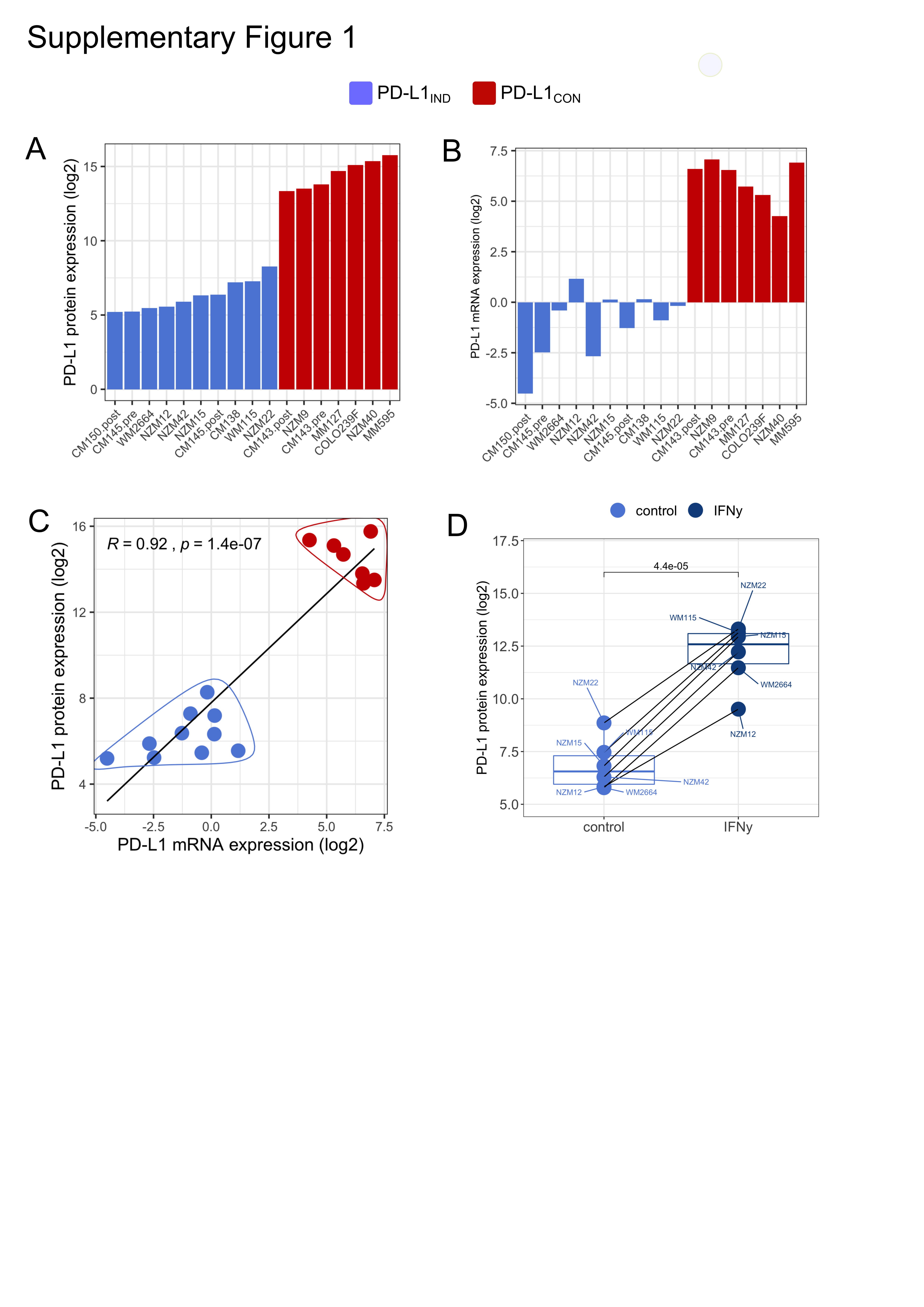

### Supplementary Figure 2

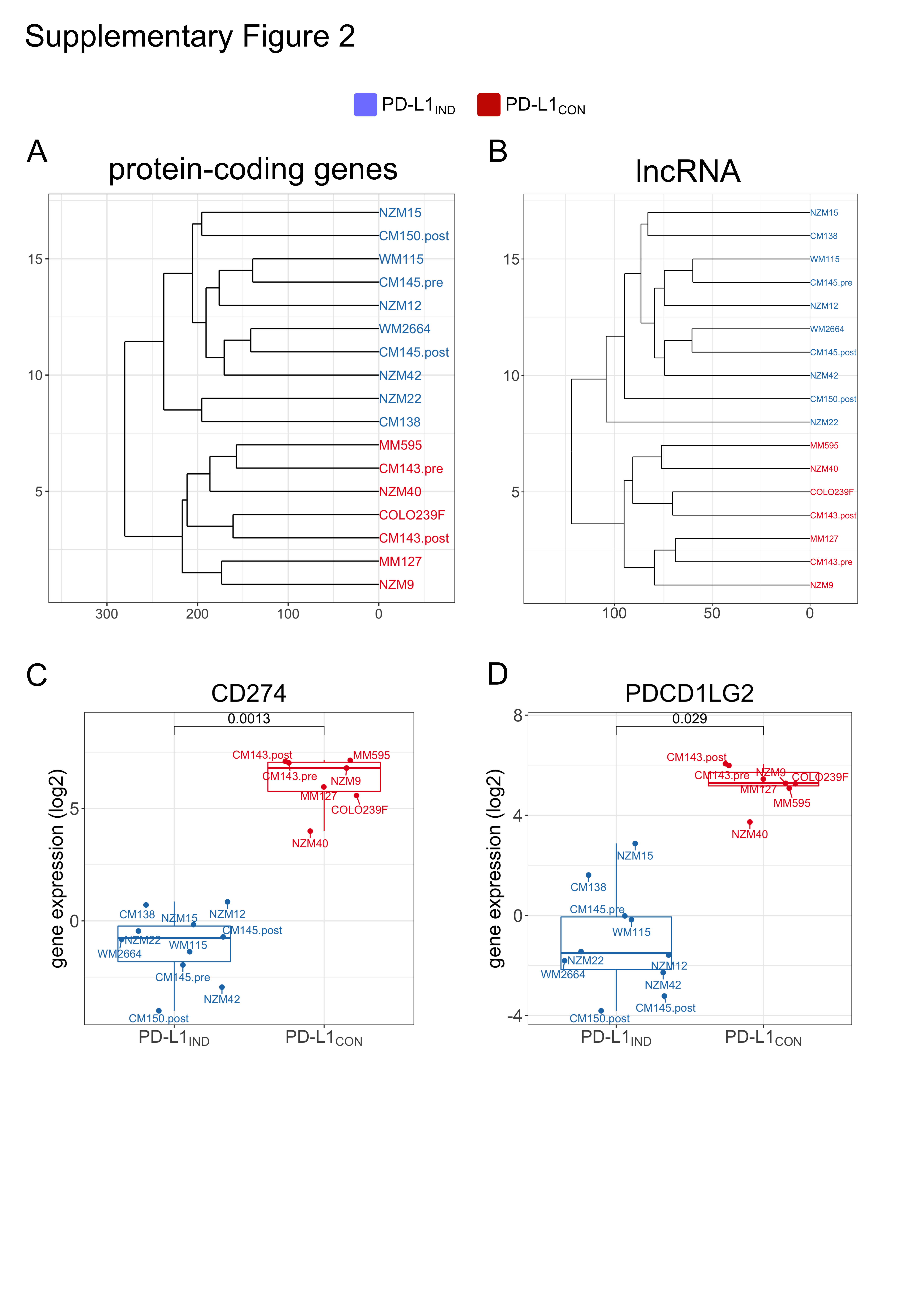

### Supplementary Figure 3

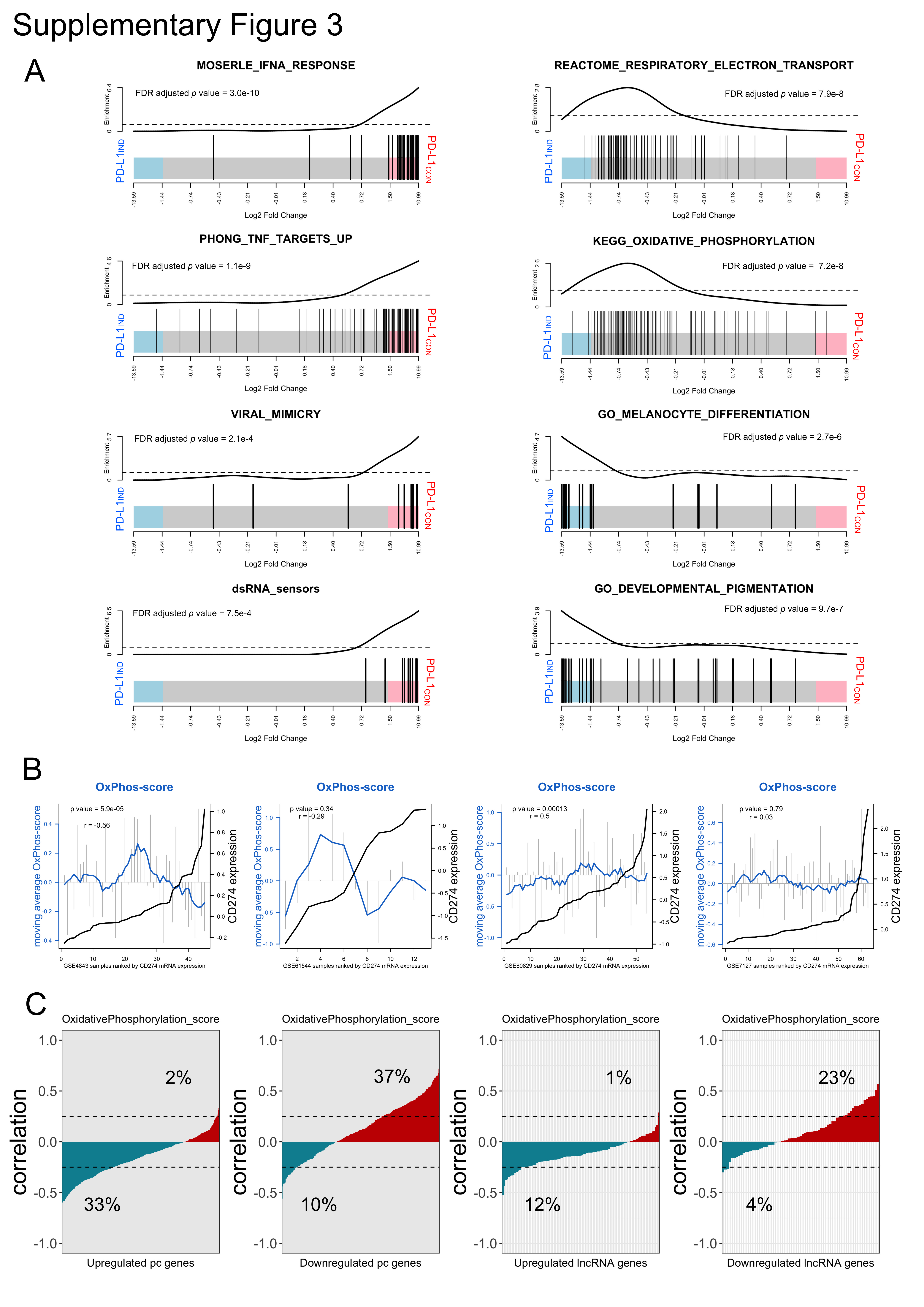

### Supplementary Figure 4

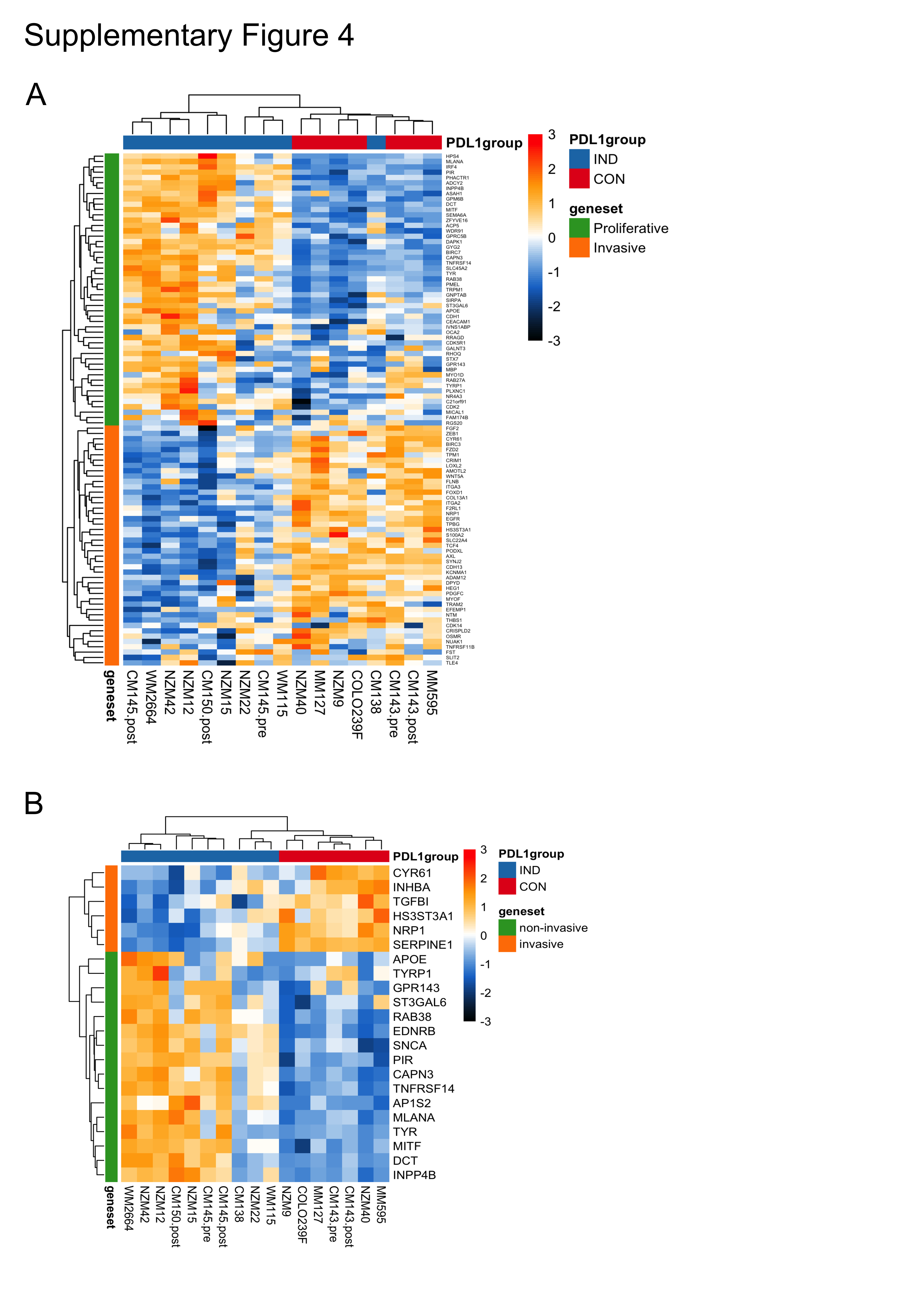

### Supplementary Figure 5

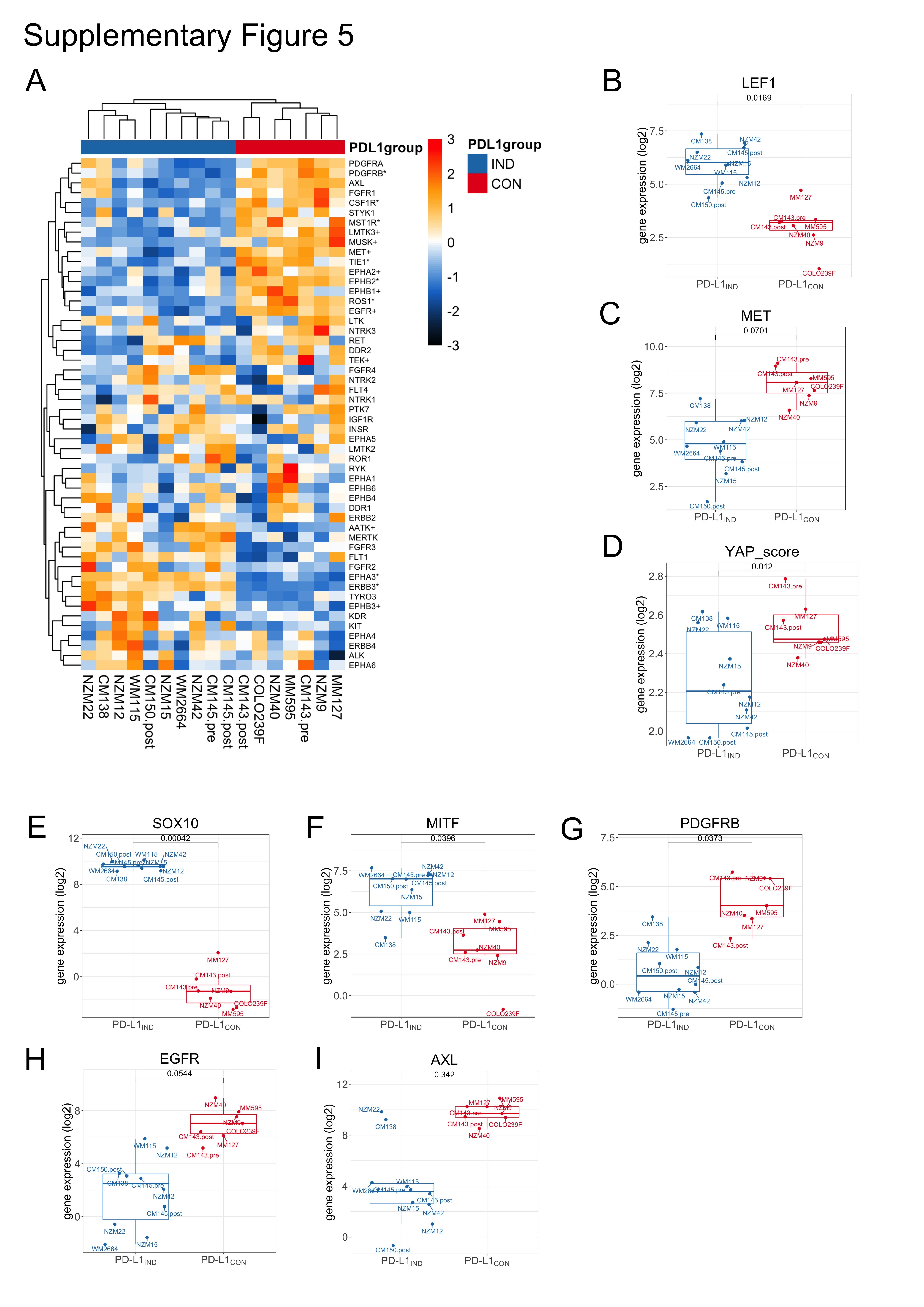

### Supplementary Figure 6

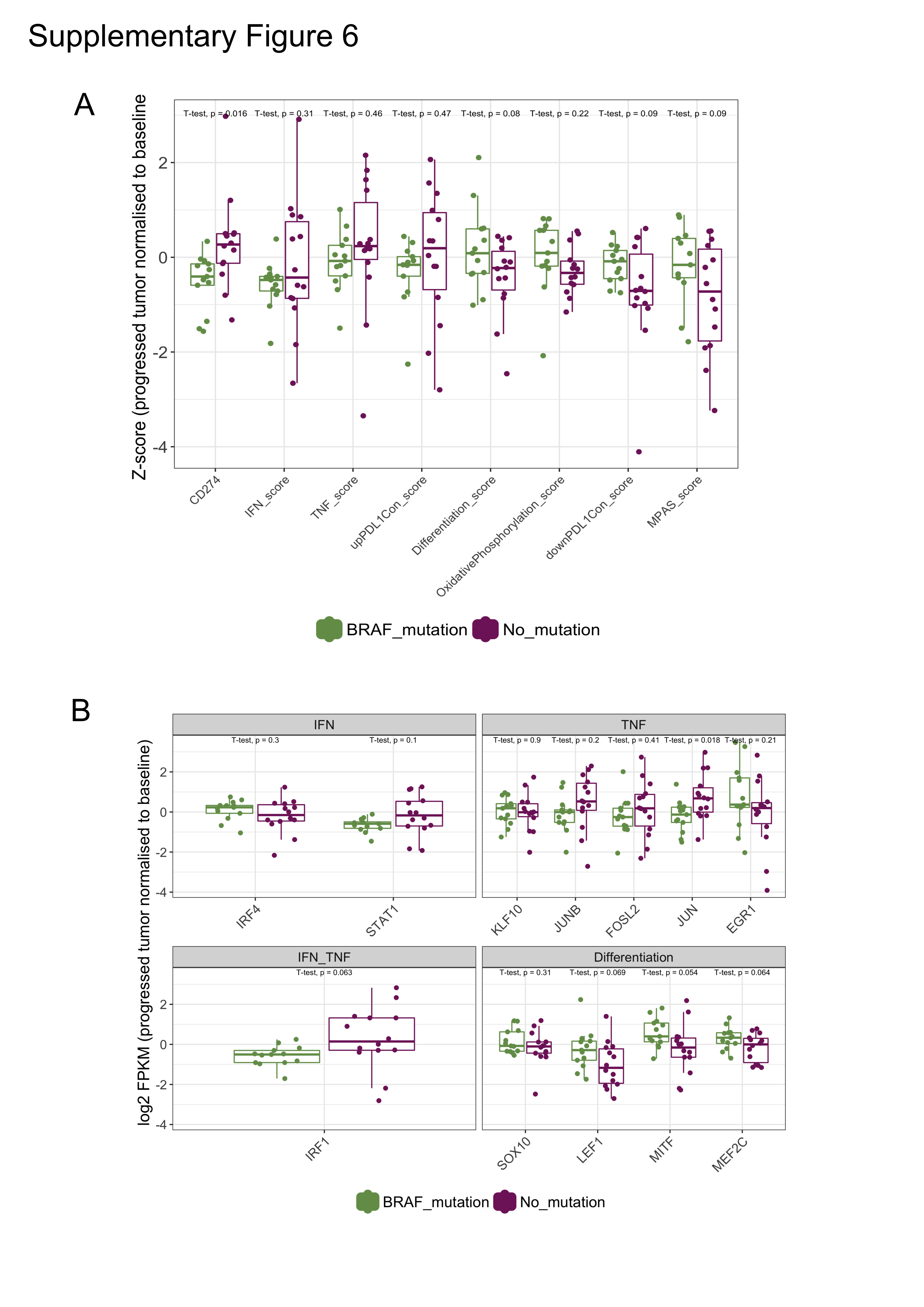

### Supplementary Table 1

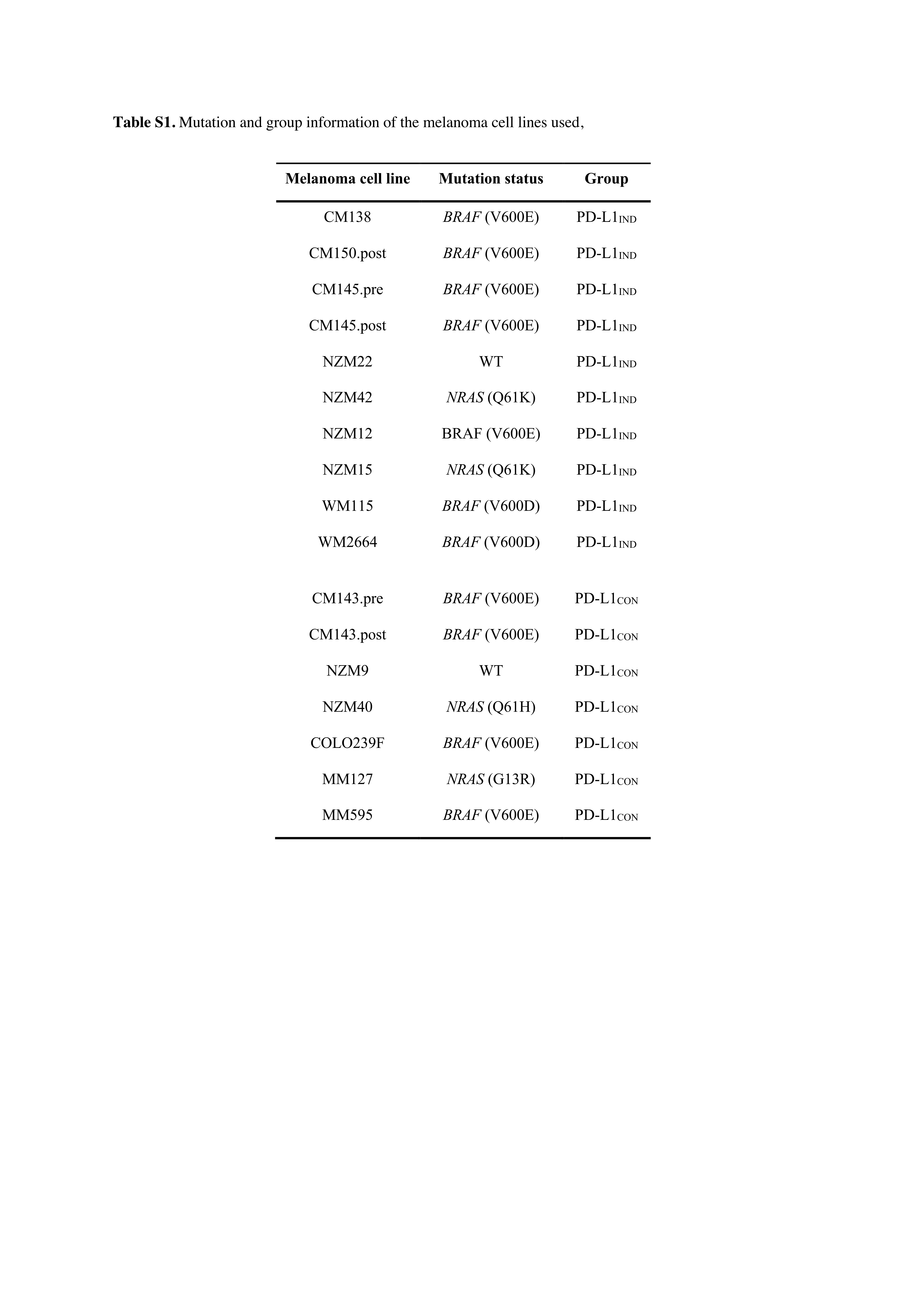
